# Gene-regulatory independent functions for insect DNA methylation

**DOI:** 10.1101/355669

**Authors:** Adam J. Bewick, Zachary Sanchez, Elizabeth C. Mckinney, Allen J. Moore, Patricia J. Moore, Robert J. Schmitz

## Abstract

The function of cytosine (DNA) methylation in insects remains unknown. Here we provide evidence for the functional role of the maintenance DNA methyltransferase 1 (*Dnmt1*) in an insect using experimental manipulation. Through RNA interference (RNAi) we successfully post-transcriptionally knocked down *Dnmt1* in ovarian tissue of the hemipteran *Oncopeltus fasciatus* (the large milkweed bug). Individuals depleted for *Dnmt1*, and subsequently DNA methylation, failed to reproduce. Eggs were inviable and declined in number, and nuclei structure of follicular epithelium was aberrant. Depletion of DNA methylation did not result in changes in gene or transposable element expression revealing an important function of DNA methylation seemingly not contingent on gene expression. Our work provides direct experimental evidence for a functional role of *Dnmt1* and DNA methylation independent of gene expression in insects.

## Introduction

DNA methylation in insects has been hypothesized to play numerous functional roles including polyphenism, diapause, longevity, and social behavior and caste differentiation [1-21]. However, these hypotheses are based primarily on correlational studies; for example, there is evidence that the dependence of social behavior and caste differentiation on DNA methylation is not absolute [11-13]. Comparative epigenomic studies have provided valuable insights into patterns of DNA methylation across most insects, but have offered limited insights into its functional significance.

In animals DNA methylation typically occurs at CG sites, which is established by DNA methyltransferase 3 (*Dnmt3*), and subsequently maintained by DNA methyltransferase 1 (*Dnmt1*). Many insects possess *Dnmt1* and *Dnmt3;* however, there is variation in presence-absence and copy number of these DNA methyltransferases, and levels and patterns of DNA methylation [12, 13]. In many insects DNA methylation is localized to moderately and constitutively expressed genes that are highly conserved between species. This has led some to the hypothesis that DNA methylation functions in transcriptional regulation [6, 15, 22, 23]. However, functional tests of DNA methyltransferases in insects are limited to the post-transcriptional knock down of *Dnmt3* in *Apis mellifera*, which is associated with modest reductions in DNA methylation, differential gene expression, and alternative splicing [24]. Given the variation in mechanisms that establish and maintain DNA methylation and limited genetic and mutant studies, the function of insect DNA methyltransferases, and DNA methylation itself, is obscure.

Here we show that DNA methylation is maintained by *Dnmt1*, is essential and is not associated with transcription in the hemipteran *Oncopeltus fasciatus* (the large milkweed bug) [25, 26]. Post-transcriptional knockdown of *Dnmt1* led to reduced egg viability, fecundity, and aberrant follicular epithelium, and thus failure to produce a successive generation. Despite finding levels of methylated CG (mCG) within coding regions reduced by 83.55%, we found no evidence for DNA methylation directly affecting transcription. Our results suggest *Dnmt1* plays an important role in reproduction in *O. fasciatus* that is mediated by a gene-regulatory independent function of DNA methylation. *Oncopeltus fasciatus* represents a fruitful model species for functional studies of DNA methylation, and continuation of studies in this system will unravel the insect epigenome and its functional consequences.

## Results

### *Dnmt1* is necessary for reproduction

To assess the function of *Dnmt1*, double-stranded RNA (dsRNA) targeting *Dnmt*1 (ds-*dnmt1*) (S1 Fig, and S1 and S2 Tables) was injected between the abdominal sternites of virgin *O. fasciatus* females. Maternal dsRNA injection has been used in *O. fasciatus* to post-transcriptionally knockdown expression in embryos developing from eggs laid by injected females [27] and is known to reduce expression of embryonically expressed genes. A control injection of double-stranded *Red* (*ds-red*) or buffer was used to confirm that there was no technical affect associated with RNAi treatment. The effects of RNAi treatment on gene expression was assessed in gut, head, thorax and ovary tissues 10 days after mating to untreated *O. fasciatus* males. The post-transcriptional knockdown reduced *dnmt1* mRNA expression in all examined tissues (Fig 1A, and S2 Fig and S2 Table). Moreover, there was little variation in expression between biological replicates in ds-*dnmt1*-injected and control individuals (Fig 1A). Taken together, treatment with ds-*dnmt1* specifically and reliably knocked down *Dnmt1* transcripts.

**Fig. 1.**
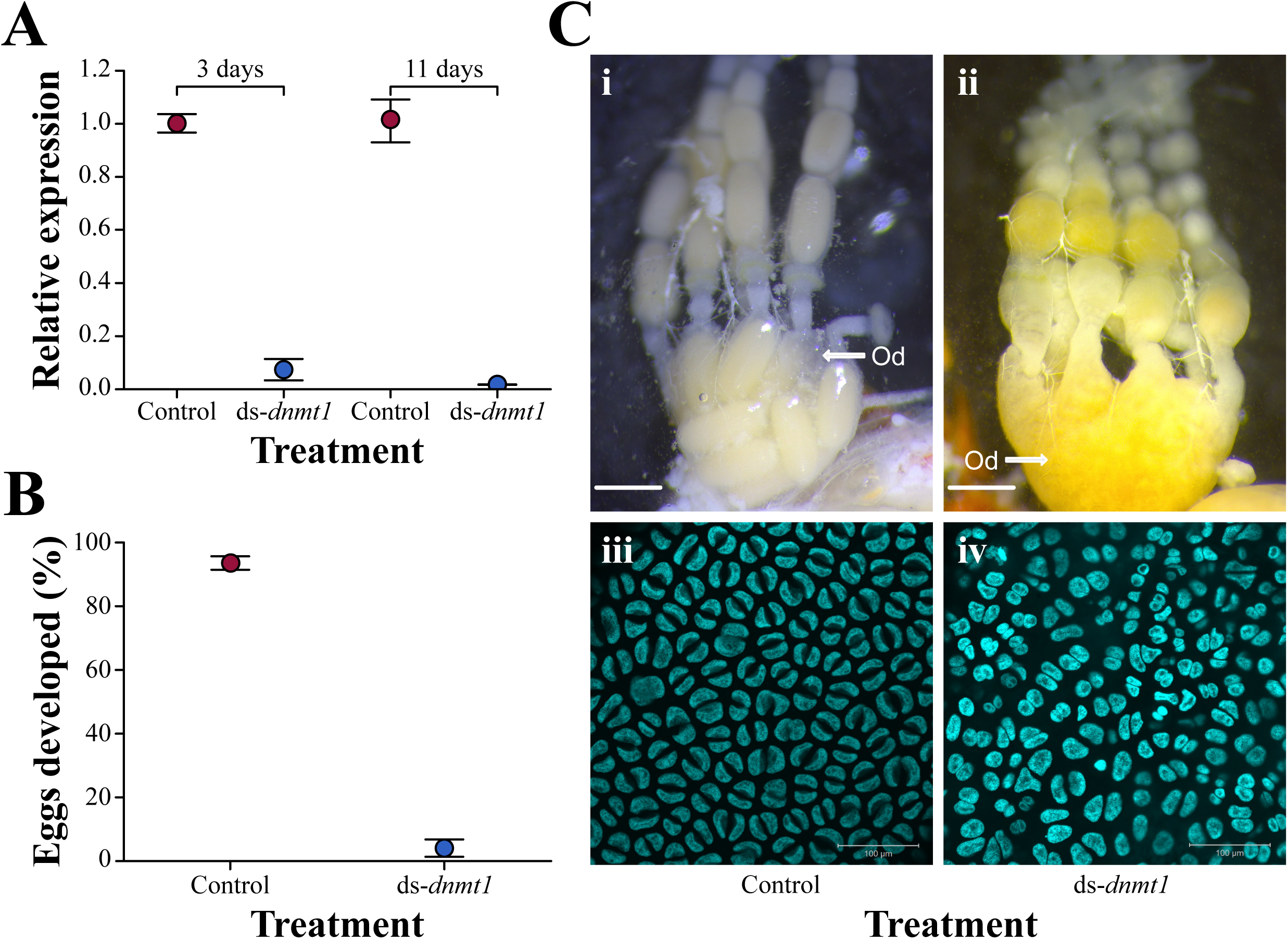
*Dnmt1* is required for reproduction in *O. fasciatus*. (A) Assessment of RNAi treatment targeting *Dnmt1* using qRT-PCR demonstrates successful reduction in transcription in ovaries compared to control. Dots indicate mean expression level, and error bars indicate standard error of the mean. (B) Parental RNAi injection with *ds*-*dnm1* significantly affected the development of eggs laid by injected females compared to eggs laid by control females. Dots indicate mean expression level, and error bars indicate standard error of the mean. (C) Whole ovaries from females removed 12-14 days post-injection (i and ii). In control females (i), mature oocytes can be seen collecting in the lateral oviduct (Od). In ds-*Dnmt1* females (ii), no mature oocytes are apparent, but the oviduct has filled with a yolk-like substance. Scale bar equals 1 mm in (i) and (ii). High magnification of the follicular epithelium surrounding a maturing oocyte (iii and iv). Nuclei are stained with Hoechst 33258 and artificially colored turquoise. Nuclei from control follicular epithelium (iii) are round and regular in shape. Nuclei from ds-*dnmt1* females (iv) are highly irregular in shape.

Post-transcriptional knockdown of *Dnmt1* affects egg development and viability. In the early stages following injection, females injected with ds-*dnmt1* did not lay fewer eggs (*F* = 2.91, *df=* 1; *p* = 1.1e-01) (S3 Fig). Although eggs laid by females injected with ds-*dnmt1* within the first 8 days post-injection looked typical, they were significantly less likely to develop than the eggs laid by control females (*χ^2^* = 8.470; *df* = 1; *p* = 4.0e-03). Furthermore, a mean of 93% of the eggs laid by control females initiated development whereas only a mean of 4% of the eggs laid by ds-*dnmt1* females initiated development (Fig 1B). Although eggs laid by the control females that initiated development were viable and hatched, the few eggs laid by ds-*dnmt1-injected* females that initiated development were not viable and failed to hatch.

Post-transcriptional knockdown of *Dnmt1* affects egg production following the first 10-day post-injection. By 10 to 12 days, females injected with ds-*dnmt1* have mainly stopped laying eggs. While ovaries dissected from control females at this stage have intact eggs in the oviduct, the oviducts of ds-*dnmt1*-injected females are either empty or filled with a mass of what appears to be yolk (Fig 1C), indicating a fault in production of a functional chorion. Analysis of ovarian cell structure indicates that knockdown of *Dnmt1* transcripts affects the cell structure of the follicular epithelium, which is responsible for production of the chorion and vitelline envelope. Nuclei of the follicular epithelium in *Dnmt1* post-transcriptional knock downed females are aberrant and fewer in number than that of the control females (Fig 1C). Therefore, both through disruption of embryonic development from eggs produced within a week of injection and cessation of egg production, reproduction is compromised following knockdown of *Dnmt1* transcripts, preventing a successive generation.

### Post-transcriptional knock down of *Dnmt1* successfully and severely reduces mCG in ovaries

The successful knockdown of *Dnmt1* transcripts in gut, head, thorax and ovaries prompted the evaluation of the consequences on DNA methylation using whole genome bisulfite sequencing (WGBS). We used a low coverage sequencing approach, which is sufficient for the detection of changes in bulk levels of DNA methylation [28]. Although *Dnmt1* mRNA expression was reduced in all tissues, DNA methylation was only reduced in ovary tissue (Fig 2A–C and S2 Fig). The reduction specifically in ovaries is likely due to the higher rate of cell division in comparison to cells in other tissues surveyed, which would facilitate the passive of loss of DNA methylation in the absence of *Dnmt1*.

**Fig. 2.**
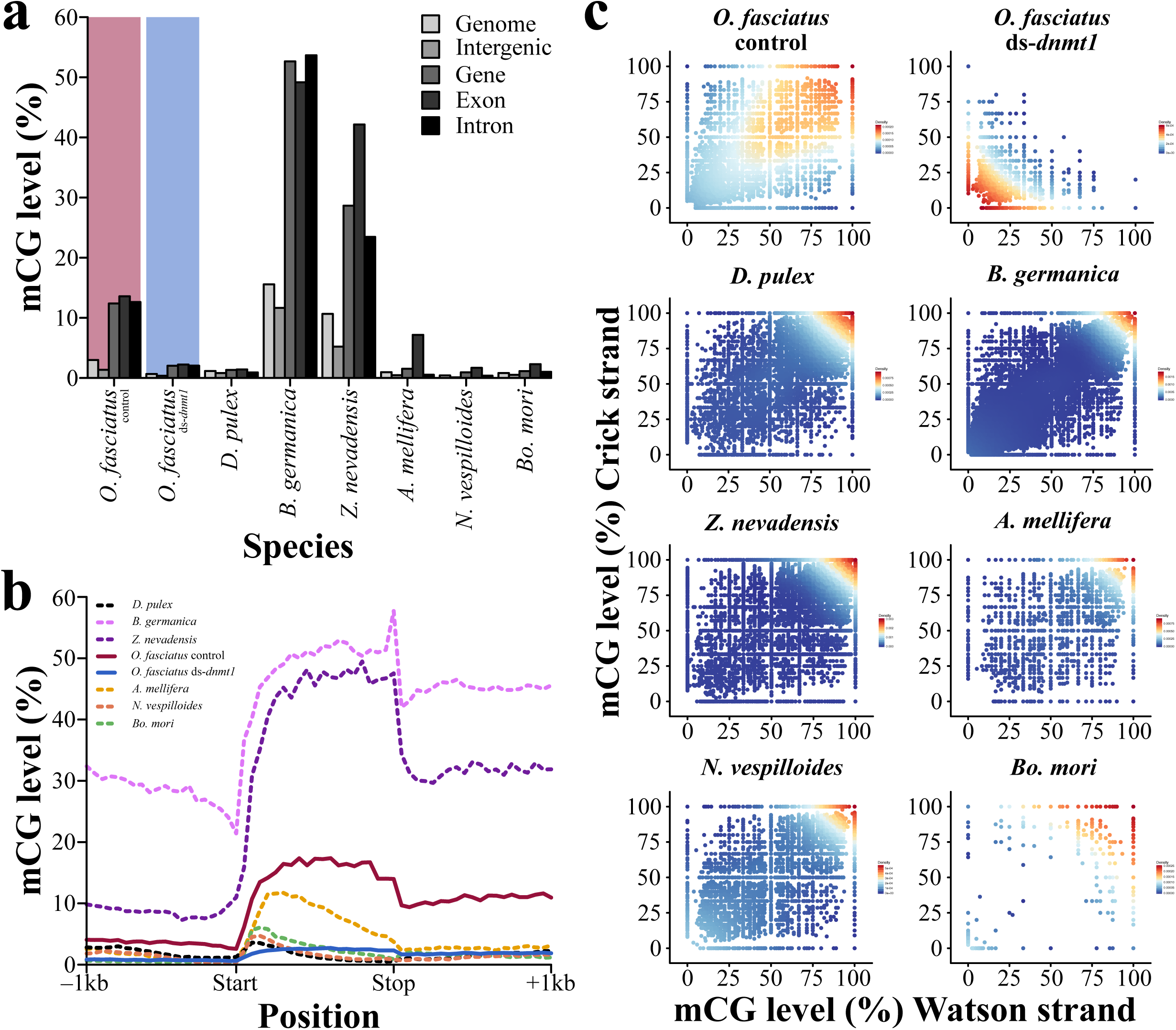
*Dnmt1* is required for mCG in *O. fasciatus*. (A) Level and genomic location of mCG between hemi– and holometabolous insects, and *D. pulex*, and RNAi treatment targeting *Dnmt1* in *O. fasciatus*. (B) Levels of mCG across gene bodies and one kilobase pairs (1kb) flanking sequence of hemi– and holometabolous insects, and *D. pulex*. Also shown are the levels of mCG across gene bodies for *O. fasciatus* ds-*dnmt1*. (C) Density plot representation of mCG for RNAi treated *O. fasciatus* and species investigated in this study.

High coverage single-base resolution DNA methylomes from ds-*dnmt1* and control ovaries were generated to understand the impact of the loss of *Dnmt1*. Greater than 75% reductions of mCG were observed across the genome, consistent with the decrease observed in the low coverage experiments (S2 Fig). The presence of symmetrical mCG – DNA methylation occurring on both DNA strands at a CpG site – is indicative of the presence of a functional *Dnmt1*. In control individuals methylation at CpGs is highly symmetrical and highly methylated. However, even though knockdown of *Dnmt1* significantly reduces methylation in ovaries, a minority of CpG sites remain symmetrically methylated (Fig 2C). This suggests that there are some cells within ovaries that have wild-type methylomes. The lack of a complete loss of mCG is expected due to the injection of fully developed individuals and hence the presence of fully methylated genomes that existed prior to the injection of dsRNA.

### Comparative epigenomic analysis of the *O. fasciatus* methylome with other insects

The *O. fasciatus* methylome revealed much higher levels of mCG throughout the genome, similar to other hemimetabolous insects, when compared to holometabolous insects (Fig 2A and B). DNA methylation of gene bodies is found in *O. fasciatus* similar to other insects that possess DNA methylation, however, the pattern of mCG within exonic regions of hemi– and holometabolous insects is distinct (Fig 2B). Higher levels of mCG are towards the 5’ end compared to the 3’ end of coding regions in holometabolous insects. This distribution resembles the most recent common ancestor of all insects (crustaceans) represented by *Daphnia pulex*. Hemimetabolous insects, which include *O. fasciatus*, have a more uniform distribution of mCG across coding regions, and higher levels towards the 3’ end of coding regions. Previous studies have demonstrated that CG-methylated genes are often conserved within insects. We identified 39.31% (N = 7,561) CG-methylated genes in *O. fasciatus* and 85.99% of these are CG-methylated in at least one other insect species investigated (S3 Table). Therefore, even though the pattern of DNA methylation within gene bodies of hemimetabolous insects is distinct compared to holometabolous insects, the targeting of specific genes is conserved. Thus, the reductions of mCG across gene bodies in ds-*dnmt1* individuals provides an apportunity to study its potential function.

### Decreased levels of mCG are not associated with transcriptome-wide changes in gene or transposable element expression in ovaries

We performed RNA-seq analysis from ovary and other tissues in ds-*dnmt1* and control individuals to better understand the relationship between gene expression and DNA methylation. No relationship between discrete mCG levels and gene expression was observed and this was consistent in other insects that we investigated (Fig 3A, and S4 Fig and S4–7 Tables). Furthermore, no correlation between continuous mCG levels and gene expression was observed (Fig 3B, and S4 Fig and S4–7 Tables). Interestingly, genome-wide relationships between DNA methylation and gene expression were not impacted by the knockdown of *Dnmt1* transcripts (Fig 3B).

**Fig. 3.**
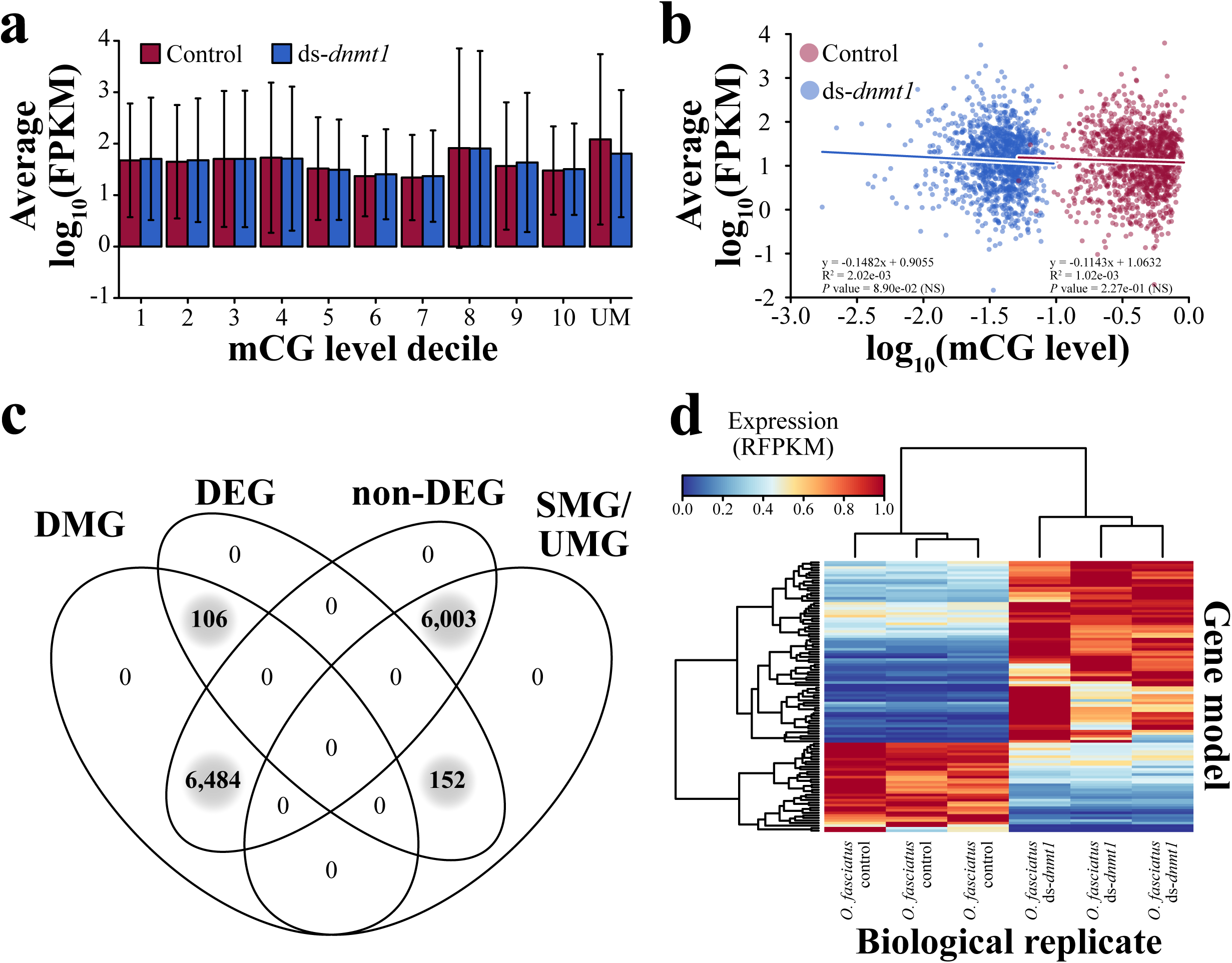
Loss of mCG in *O. fasciatus* ovaries has a limited effect on transcription. (A) Gene expression level for deciles of increasing mCG (1–10) and unmethylated genes (UM). Error bars represent 95% confidence interval of the mean. (B) Regression of gene expression against a continuous measure of mCG with > 0 FPKM for the same set of genes that are CG-methylated in *O. fasciatus* control, but unmethylated in *ds-dnmt1*. Raw *p* values are provided for each regression, and significance or non-significance (NS) is indicated in brackets following Bonferroni correction. (C) Combinational overlap of genes that are differential CG-methylated and expressed, and similarly CG-methylated and expressed between *O. fasciatus ds-dnmt1* and control. Gene groups: Differentially Methylated Gene (DMG), Differentially Expressed Gene (DEG), Similarly Methylated Gene (SMG)/UnMethylated Genes (UMG), and non-Differentially Expressed Gene (non-DEG). (D) A heatmap showing gene expression changes for genes that are differentially CG-methylated between *O. fasciatus* ds-*dnmt1* and control. Expression was standardized by the highest value per gene per biological replicate to produce a Relative Fragments Per Kilobase of transcript per Million (RFPKM) value. RFPKM were clustered using a hierarchical clustering method.

A maximum of 264 differentially expressed genes (DEG) were observed between *O. fasciatus* ds-*dnmt1* and control ovaries (S5 Table). *Dnmt1* was down-regulated in all ds-*dnmt1* samples and no detection of the *de novo* DNA methyltransferase *Dnmt3* was observed. Additionally, no DEGs are observed between *O. fasciatus* ds-*dnmt1* and control for gut, head and thorax (S8 Table). Of genes with differences in DNA methylation between ds-*dnmt1* and control ovaries (N = 6,590), the majority (N = 6,484; 98.39%) had no changes to gene expression (Fig 3C, and S4 and S5 Tables). Furthermore, mCG was always reduced in ds-*dnmt1* compared to control ovaries in the 6,484 gene set. Despite genes being unmethylated (UM) in ds-*dnmt1* ovaries, changes to gene expression occurred in both directions for the 106 differentially DNA methylated and expressed genes (Fig 3D). Even for genes that by definition are unmethylated (N = 5,982) or similarly CG-methylated (N = 21) in ds-*dnmt1* and control, a similar proportion (N = 6,003; 97.53%) was observed to have no difference in gene expression. The lack of an association between DNA methylation and gene expression is further supported when applying more stringent thresholds to the definition of CG-methylated and unmethylated genes (S5 Fig). Our observations support a trivial role of DNA methylation in gene expression if any at all.

The function of DNA methylation in *O. fasciatus* might lie within genome defense – the transcriptional regulation of repetitive DNA and transposons – as its genome is composed of a nontrivial amount of repetive DNA and transposons (6.21%) (S9 Table). However, as in genes, reduction of mCG in ds-*dnmt1* compared to control ovaries was found across the bodies of the TEs, and were not associated with a genome wide increase in expression (Fig 4A–C, and S6 Fig and S10 Table). This further supports a trivial role of DNA methylation in transcriptional regulation of loci.

**Fig. 4.**
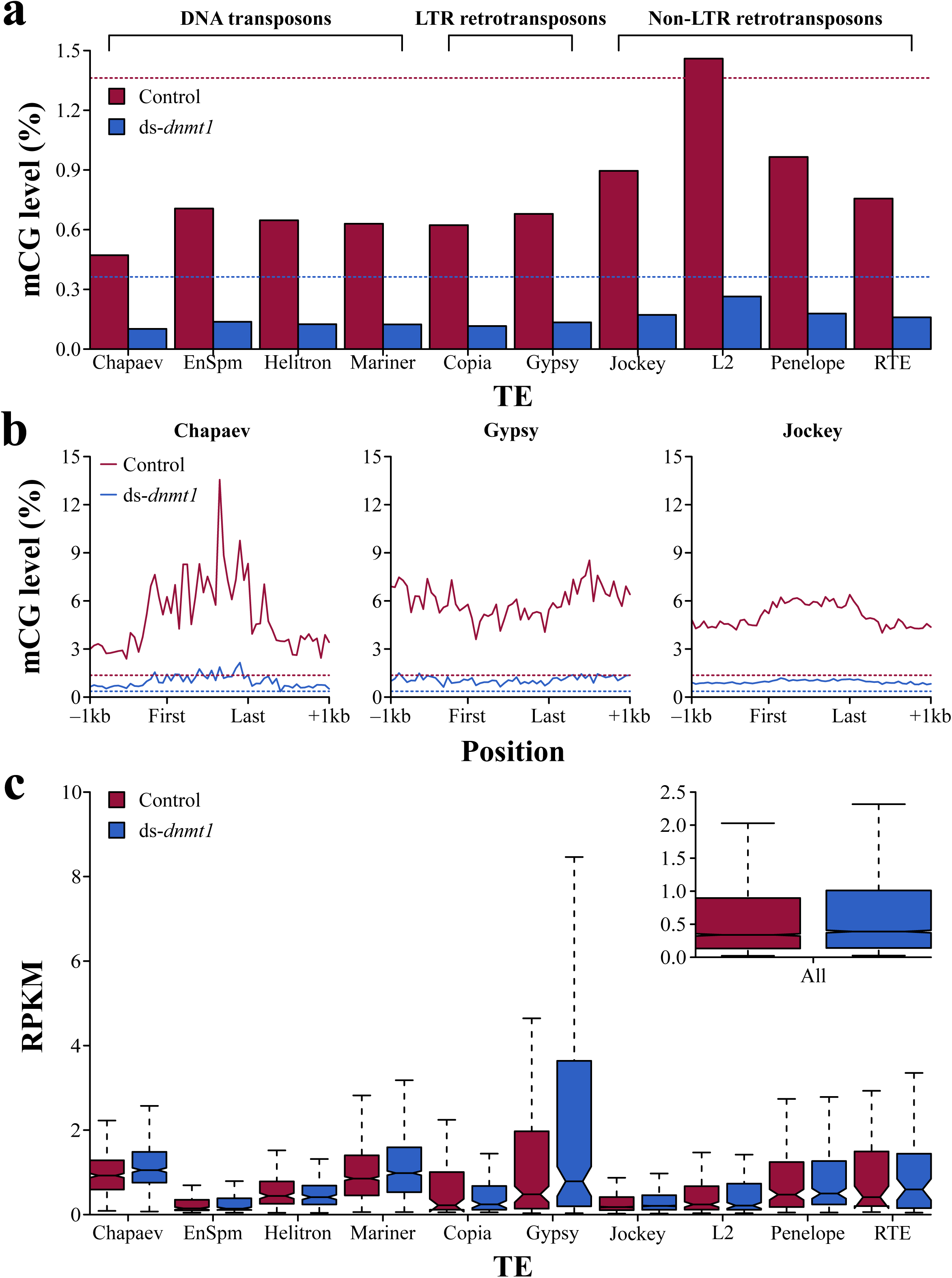
Large-scale reactivation of TEs did not follow severe reductions of mCG in *O. fasciatus* ovaries. (A) CG methylation levels for the top ten most abundant TEs of ≥ 500 bp in the *O. fasciatus* genome. The dashed lines correspond to the intergenic mCG level of *O. fasciatus* ds-*dnmt1* and control. (B) Levels of mCG across the bodies and 1kb flanking sequence of TEs for a single representative of DNA transposons (Chapaev), LTR retrotransposons (Gypsy), and non-LTR retrotransposons (Jockey). (C) Expression quantified as RPKM for the top ten most abundant TEs of ≥ 500 bp. The subset presents the RPKM distribution when all TEs are considered within ds-*dnmt1* and control ovaries.

## Discussion

DNA methylation has been implicated to play numerous roles in insects [1–21], however, its exact function remains uncertain. In this study we observed that post-transcriptional knockdown of *Dnmt1* negatively impacts reproduction through a decrease in egg development and egg production in *O. fasciatus*.

Post-transcriptional knockdown of *Dnmt1* transcripts impacted the maternal somatic gonad and egg maturation. The follicular epithelial cells undergo multiple rounds of mitosis during oogenesis [29]. Oocytes that were in the early stages of maturation would be the most effected, as their follicular epithelium would passively lose DNA methylation with each successive round of mitotis. Loss was not compensated by the *de novo* DNA methyltransferase *Dnmt3* as expression was not observed in ovaries. The loss of function of follicular epithelial cells, which includes production of the chorion and vitelline membrane, could explain the loss of oocyte integrity.

The knockdown of *Dnmt1* transcripts in *O. fasciatus* ovaries subsequently did not lead to transcriptome-wide changes in gene expression. No correlation between mCG and gene expression in *O. fasciatus* ovaries and other insects was observed (Fig 3 and S4 Fig) [6, 13]. Hence, the importance of mCG in *O. fasciatus* might lie within the orchestration of genomic structure during cell replication, gametogenesis, or cell types not examined in our study.

DNA methylation might be required for proper mitosis, and the segration of sister chromatids into their respective daughter cells, as there does appear to be aberrant nuclei structure in ds-*dnmt1* compared to control ovarian cells (Fig. 1C). This phenotype could be the result of epigenomic defects in scaffold/matrix attachment regions (S/MAR), as some correlate with origins of replication [30]. This phenotype could be exacerbated by multiple mitotic divisions of follicular epithelial cells during oogenesis, and/or the presence of holocentric chromosomes [31]. Future work describing the epigenomic contributions to cellular defects are now possible to study in this newly emerging and tractable model species, *O. fasciatus*.

We suggest that DNA methylation is more important for genome structure, integrity or other cellular processes than it is for somatic expression in *O. fasciatus*. Although it is possible that DNA methylation is important for expression control in a rare cell type that was not examined in our study, we instead propose that regulation of expression is not likely the primary role in insects. Investigating the importance of DNA methylation prior to the onset of meiosis and throughout embryo development is a fundamental next-step. *Oncopeltus fasciatus* represents a fruitful model species for functional studies of DNA methylation, and continuation of studies in this system will unravel the insect epigenome and its functional consequences.

## Materials and Methods

### Phylogenetic analysis

A subset of DNA methyltransferase (*Dnmt*) 1, 2, and 3 sequences was obtained from [16] for phylogenetic analysis. The subset included only insect species with available MethylC-seq data, and species representatives from the dipteran suborders Brachycera and Nematocera: *Acyrthosiphon pisum*, *Aedes aegypti*, *Aedes albopictus*, *Anopheles gambiae*, *Apis mellifera*, *Bombyx mori*, *Camponotus floridanus*, *Copidosoma floridanum*, *Culex pipiens quinquefasciatus*, *Drosophila melanogaster*, *Harpegnathos saltator*, *Microplitis demolitor*, *Nasonia vitripennis*, *Nicrophorus vespilloides*, *O. fasciatus*, *Ooceraea* (*Cerapachys*) *biroi*, *Polistes canadensis*, *Polistes dominula*, *Solenopsis invicta*, *Tribolium castaneum*, and *Zootermopsis nevadensis*. DNA methyltransferases were reassessed in *O. fasciatus* by using InterProScan v5.23-62.0 [32] to identify annotated proteins with a C-5 cytosine-specific DNA methylase domain (PF00145). Sequence identifiers are located in S1 Fig. Full-length protein sequences were aligned using PASTA v1.6.4 [33], and manually trimmed of divergent, non-homologous sequence in Mesquite v3.2 [34]. Full, aligned, and aligned and trimmed sequence alignments, and parenthetical phylogenetic tree are located in S1 Data. Phylogenetic relationship among *Dnmt* sequences was estimated using BEAST v2.3.2 [35] with a Blosum62+Γ model of amino acid substitution. A Markov Chain Monte Carlo (MCMC) was ran until stationarity and convergence was reached (10,000,000 iterations), and a burnin of 1,000,000 was used prior to summarizing the posterior distribution of tree topologies. A consensus tree was generated using TreeAnnotator v2.3.2, visualized in FigTree v1.4.2 (http://tree.bio.ed.ac.uk/software/figtree/) and exported for stylization in Affinity Designer v1.5.1 (https://affinity.serif.com/en-us/).

### PCR confirmation for the presence of a single *Dnmt1* ortholog in *O. fasciatus*

To determine if OFAS015351 and OFAS018396 were two parts of a single *Dnmt1* ortholog, we designed one sense primer at the 3’ end of OFAS015351 (Of_DMNT1-1_3603S; S1 Table) and two antisense primers at the 3’ end of OFAS018396 (Of_DNMT1-2_424A and Of_DNMT1-2_465A; S1 Table). A fourth sense primer (Of_DNMT1-2_1S; S1 Table) was designed at the 5’ end of OFAS018396 to confirm the size and sequence of this possibly truncated gene annotation.

Polymerase Chain Reaction (PCR) with primer combinations Of_DMNT1-1_3603S–Of_DNMT1-2_424A, Of_DMNT1-1_3603S–Of_DNMT1-2_465A, and Of_DNMT1-2_1S–Of_DNMT1-2_424A was performed using Q5 Polymerase (New England BioLabs, Ipswich, MA) per manufactures instructions. Thermacycler conditions were 98°C for 15 seconds (s) (denaturing), 60°C for 30 s (annealing), and 72°C for 30 s (extension), and repeated for 40 cycles. The PCR products were then purified using QIAquick PCR Purification Kit (Qiagen, Venlo, The Netherlands), and sequenced at the Georgia Genomics and Bioinformatics Core (Athens, GA).

### Animal culture

*Oncopeltus fasciatus* cultures were originally purchased from Carolina Biologicals (Burlington, NC). Mass colonies were maintained in incubators under a 12 h:12 h light/dark cycle at 27°C. Colonies and individual experimental animals were fed organic raw sunflower seeds and provided with *ad libitum* deionized water. Late instar nymphs were separated from the mass colonies and housed under the same conditions. Nymph colonies were checked daily for newly emerged adults. Adults were separated by sex and kept with food and water for 7–10 days until females reached sexual maturity.

### Parental RNAi

Template for the *in vitro* transcription of reactions was prepared from a PCR reaction in which T7 phage promoter sequences were added to the gene-specific *Dnmt1* primers [36]. For our control sequence, we used the red fluorescent protein (*Red*) sequence used in previous parental RNAi experiments in *O. fasciatus* [36] or buffer. Primer sequences can be found in S2 Table. Sense and anti-sense RNA was synthesized in a single reaction using the Ambion MEGAscript kit (ThermoFisher Sci, Waltham, MA). After purification, the double-stranded RNA (dsRNA) concentration was adjusted to 2 μg/μL in injection buffer (5 mM KCl, 0.1 mM NaH2PO4) [36]. Females were injected with 5 μL of dsRNA between the abdominal sternites using an insulin syringe. Following injection, females were paired with an un-injected male to stimulate oviposition and fertilize eggs. Individual females with their mate were housed in petri dishes with sunflower seeds, water and cotton wool as an oviposition site. The parental RNAi protocol has been reported to result in 100% penetrance by the third clutch of eggs [36] and this was also our experience.

### Reproductive phenotype screening and analysis

Eggs were collected between days 4–10 post-injection and assessed for development. *Oncopeltus fasciatus* embryos change from a creamy white color to orange as they develop, which indicates viability. Thus, color change is a useful tool for assessing healthy development. We examined the number of developing eggs at 5 days post-oviposition, at which point viable eggs are clearly distinguishable from inviable eggs, as well as hatching rate of eggs from females injected with double-stranded *Dnmt1* (ds-*dnmt1*) and double-stranded *Red* (ds-*red*). The data for the number of eggs laid was normally distributed so differences among RNAi treated groups were tested using analysis of variance (ANOVA). The development data, however, was not normally distributed and consisted of binary states (developed and not developed), and so differences among treatment groups were analyzed with a Generalized Linear Model (GLM) using a Poisson distribution.

A second set of *O. fasciatus* females were injected in the same manner as described in **Parental RNAi** to assess post-transcriptional knockdown of *Dnmt1* on ovarian structure. *Oncopeltus fasciatus* females were dissected 10 days after injection. By 10 days post-injection *O. fasciatus* females are beginning to stop laying recognizable eggs. Ovaries were removed from *O. fasciatus* females and placed in 1× PBS. Whole ovaries were imaged with a Leica DFC295 stereomicroscope using Leica Application Suite morphometric software (LAS V4.1; Leica, Wetzlar, Germany). Dissected ovaries were fixed within 15 minutes of dissection in 4% formaldehyde in 1× PBS for 25 minutes. Fixed ovaries were stained with Hoechst 33342 (Sigma Aldrich) at 0.5 µg/ml. The stained ovarioles were imaged using a Zeiss LSM 710 Confocal Microscope (Zeiss) at the University of Georgia Biomedical Microscopy Core.

### Quantitative RT-PCR

To assess the effectiveness of post-transcriptional knockdown of *Dnmt1*, females were dissected 11 days post-injection. Ovaries were removed from each female, flash frozen in liquid nitrogen and stored at −80°C until processing. Total RNA (and DNA) was extracted from a single ovary per female using a Qiagen Allprep DNA/RNA Mini Kit (Qiagen, Venlo, The Netherlands) per manufacturer’s instructions. Complementary DNA (cDNA) was synthesized from 500 ng RNA with qScript cDNA SuperMix (Quanta Biosciences, Gaithersburg, MD).

Expression level of *Dnmt1* was quantified by quantitative real-time PCR (qRT-PCR). Primers were designed for *Dnmt1* using the *O. fasciatus* genome as a reference [37]. Actin and GAPDH were used as endogenous reference genes. Primer sequences can be found in Extended Data Table 2. We used Roche LightCycler 480 SYBR Green Master Mix with a Roche LightCycler 480 (Roche Applied Science, Indianapolis, IN) for qRT-PCR. All samples were run with 3 technical replicates using 10 μL reactions using the manufacturer’s recommended protocol. Primer efficiency calculations, genomic contamination testing and endogenous control gene selection were performed as described by [37]. We used the ΔΔCT method [38] to examine differences in expression between control and ds-*dnmt1* injected females [37].

### Whole-Genome Bisulfite Sequencing (WGBS) and analysis of cytosine (DNA) methylation

MethylC-seq libraries for an *O. fasciatus* ds-*dnmt1* and control individual were prepared according to the protocol described in [39] using genomic DNA extracted from ovaries (see **Materials and Methods** section **Quantitative RT-PCR**). Libraries were single-end 75 bp sequenced on an Illumina NextSeq500 machine [40, 41]. Unmethylated lambda phage DNA was used to as a control for sodium bisulfite conversion, and an error rate of ~0.05% was estimated. *Oncopeltus fasciatus* ds-*dnmt1* and control were sequenced to a depth of ~18× and ~21×, which corresponded to an actual mapped coverage of ~9× and ~11×, respectively. Additionally, low pass (< 1×) WGBS from gut, head, ovary, and thorax was performed for three control and ds-*dnmt1* biological replicates. *Blattella germanica* was additionally sequenced to generate equal numbers of hemi– and holometabolous insects investigated in this study. However, DNA was extracted from whole-body minus gastrointestinal tract. MethylC-seq libraries were prepared and sequenced identically to *O. fasciatus*. An error rate of ~0.14% was estimated from unmethylated lambda phage DNA. *Blattella germanica* was sequenced to a depth of ~8×, which corresponded to an actual mapped coverage of ~5×. WGBS data for *O. fasciatus* and *B. germanica* can be found on Gene Expression Omnibus (GEO) under accession GSE109199. Previously published WGBS data for *A. mellifera* [42], *Bo. mori* [17], *Daphnia pulex* [43], *N. vespilloides* [19], and *Z. nevadensis* [15] were downloaded from the Short Read Archive (SRA) using accessions SRR445803–4, SRR027157–9, SRR1552830, SRR2017555, and SRR3139749, respectively. Thus, DNA methylation was investigated for six insects from six different orders spread evenly across developmental groups, and a crustacean outgroup. WGBS data was aligned to each species respective genome assembly using the methylpy pipeline [44]. In brief, reads were trimmed of sequencing adapters using Cutadapt v1.9 [45], and then mapped to both a converted forward strand (cytosines to thymines) and converted reverse strand (guanines to adenines) using bowtie v1.1.1 [46]. Reads that mapped to multiple locations, and clonal reads were removed.

Weighted DNA methylation was calculated for CG sites by dividing the total number of aligned methylated reads by the total number of methylated plus unmethylated reads [47]. For genic metaplots, the gene body (start to stop codon), 1000 base pairs (bp) upstream, and 1000 bp downstream was divided into 20 windows proportional windows based on sequence length (bp). Weighted DNA methylation was calculated for each window and then plotted in R v3.2.4 (https://www.r-project.org/). CG sequence context enrichment for each gene was determined through a binomial test followed by Benjamini-Hochberg false discovery rate [48, 49]. A background mCG level was determined from all coding sequence, which was used as a threshold in determining significance with a False Discovery Rate (FDR) correction. Genes were classified as CG-methylated if they had reads mapping to at least 20 reads mapping to 20 CG sites and a *q* < 0.05. Using a binomial test can lead to false-negatives – highly CG-methylated genes that are classified as unmethylated (UM) – due to a low number of statistically CG-methylated sites (S4 Table). Genes classified as unmethylated, but had a mCG level greater than the lowest CG-methylated gene were dropped from future analyses.

### Ortholog identification

Best BLASTp hit (arguments: -max_hsps 1 –max_target_seqs 1 evalue 1e-03) was used to identify orthologs between *O. fasciatus* and other insect species investigated.

### RNA-seq and differential expression analysis

RNA-seq libraries for RNA extracted from ovaries of three biological *O. fasciatus* ds-*dnmt1* and control replicates at 11 days post-injection were constructed using Illumina TruSeq Stranded RNA LT Kit (Illumina, San Diego, CA) following the manufacturer’s instructions with limited modifications. RNA from ovaries of an additional three biological *O. fasciatus* ds-*dnmt1* and control replicates, and three biological *O. fasciatus* ds-*dnmt1* and control replicates from gut, head, and thorax were extracted. The starting quantity of total RNA was adjusted to 1.3 µg, and all volumes were reduced to a third of the described quantity. Libraries were single-end 75 bp sequenced on an Illumina NextSeq500 machine. RNA-seq data for *O. fasciatus* ds-*dnmt1* and control can be found on GEO under accession GSE109199. Previously published RNA-seq data for *A. mellifera* [50], and *Z. nevadensis* (15) were downloaded from the SRA using accessions SRR2954345, and SRR3139740, respectively.

Raw RNA-seq FASTQ reads were trimmed for adapters and preprocessed to remove low-quality reads using Trimmomatic v0.33 (arguments: LEADING:10 TRAILING:10 MINLEN:30) [51] prior to mapping to the *O. fasciatus* v1.1 reference genome assembly. Reads were mapped using TopHat v2.1.1 [52] supplied with a reference General Features File (GFF) to the *O. fasciatus* v1.1 reference genome assembly [26], and with the following arguments: -I 20000 --library-type fr-firststrand --b2-very-sensitive.

Differentially expressed genes (DEGs) between ds-*dnmt1* and control libraries were determined using edgeR v3.20.1 [53] implemented in R v3.2.4 (https://www.r-project.org/). Genes were retained for DEG analysis if they possessed a Counts Per Million (CPM) ≥ 1 in at least ≥ 2 libraries. Significance was determined using the glmQLFTest function, which uses empirical Bayes quasi-likelihood F-tests. Parameter settings were determined following best practices for DEG analysis as described by [54]. Gene expression metrics for *A. mellifera*, *Z. nevadensis*, and *O. fasciatus* ds-*dnmt1* and control are located in S5–8 Tables, respectively.

### Gene Ontology (GO) annotation and enrichment

GO terms were assigned to *O. fasciatus* v1.1 gene set [26] through combining annotations from Blast2Go PRO v4.1.9 [55], InterProScan v5.23-62.0 (arguments: -goterms -iprlookup -appl CDD,Pfam) [32], and through sequence homology to *D. melanogaster* using BLASTp (arguments: -evalue 1.0e-03 -max_target_seqs 1 -max_hsps 1). 11,105/19,615 gene models were associated with at least one GO term and a total of 8,190 distinct GO identifiers were mapped. GO terms are found in S11 Table. Enriched GO terms in gene groups were evaluated using topGO v2.30.0 [56] implemented in R v3.2.4 (https://www.r-project.org/), and significance (*p* < 0.05) of terms was assessed using Fisher’s exact test with a weighted algorithm. Gene groups were contrasted to all *O. fasciatus* genes associated with GO terms (S12 Table).

### Transposable element (TE) annotation and expression

TEs were identified using RepeatMasker v4.0.5 (http://www.repeatmasker.org) provided with the invertebrate repeat library from Repbase (http://www.girinst.org/repbase/) (arguments: -lib <Repbase invertebrate library> -no_is -engine wublast -a -inv -x -gff. Following RepeatMasker, neighboring TEs of the same type were collapsed into a single locus within the outputted GFF. The unmodified GFF is located in S9 Table.

To quantify expression from TEs RNA-seq libraries from *O. fasciatus* ds-*dnmt1* and control were independently combined and mapped to the *O. fasciatus* v1.1 reference genome assembly [26] using bowtie2 v2.2.9 [57] with the following arguments: -- sensitive. Mapped reads overlapping with the top ten most abundant TEs of ≥ 500 bp in length were identified using the *intersect* command in BEDTools suite v 2.26.0 [58]. TE expression is quantified as Reads Per Kilobase per Million mapped reads (RPKM) for each intersected TE type by counting the number reads and dividing by the mapped library read number in millions. Significance in expression of TEs between ds-*dnmt1* and control tissues was assessed using the Mann-Whitney test with the alternative hypothesis set to “greater” in R v3.2.4 (https://www.r-project.org/).

## Acknowledgements

We thank Sam Arsenault (University of Georgia [UGA]), and Mariana Monteiro and Stefan Götz (Blast2GO Team) for assistance with Blast2GO analyses, Nick Rohr and Tina Ethridge for MethylC-Seq and RNA-seq library preparation, Brigitte Hofmeister (UGA) for bioinformatics advice, Mary Goll (UGA) for comments, Kelly Dawe (UGA) for feedback on cellular phenotypes, Muthugapatti Kandasamy of the Biomedical Microscopy Core (UGA) for technical assistance with confocal microscopy, and Cassandra Extavour (Harvard) for providing technical advice and *Red* plasmids for RNAi. We also thank the Georgia Advanced Computing Resource Center (GACRC), Georgia Genomics and Bioinformatics Core (GGBC) and the Office of Research at the University of Georgia. This work was funded by the National Science Foundation (NSF) IOS-1354358 (AJM), and UGA CAES Undergraduate Research Award (ZS). RJS is a Pew Scholar in the Biomedical Sciences, supported by The Pew Charitable Trusts.

## Supporting Information

**S1 Fig. Identification of *O. fasciatus* DNA methyltransferases.** (A) Phylogenetic relationship of DNA methyltransferases identified the *de novo* (*Dnmt3*) and maintenance (*Dnmt1*) DNA methyltransferase in *O. fasciatus*. Node support with ≤0.5 posterior probability is indicated – other nodes are ≥0.95. Branch lengths are in amino acid substitutions per site. Species names are represented as abbreviations: Acy. pis.: *Acyrthosiphon pisum*, Aed. aeg.: *Aedes aegypti*, Aed. alb.: *Aedes albopictus*, Ano. gam.: *Anopheles gambiae*, Api. mel.: *A. mellifera*, Bom. mor.: *Bo. mori*, Cam. flo.: *Camponotus floridanus*, Cop. flo.: *Copidosoma floridanum*, Cul. qui.: *Culex pipiens quinquefasciatus*, Dro. mel.: *Drosophila melanogaster*, Har. sal.: *Harpegnathos saltator*, Mic. dem.: *Microplitis demolitor*, Nas. vit.: *Nasonia vitripennis*, Nic. ves.: *Nicrophorus vespilloides*, Onc. fas.: *O. fasciatus*, Cer. bir.: *Ooceraea* (*Cerapachys*) *biroi*, Pol. can.: *Polistes canadensis*, Pol. dom.: *Polistes dominula*, Sol. inv.: *Solenopsis invicta*, Tri. cas.: *Tribolium castaneum*, and Zoo. nev.: *Z. nevadensis*. (B) A to scale representation of *Dnmt1* and protein domains identified in *O. fasciatus* and *M. musculus*.

**S2 Fig. DNA methylation consequences following post-transcriptional knockdown of *Dnmt1* are restricted to ovaries.** (A) Assessment of RNAi treatment targeting *Dnmt1* using qRT-PCR demonstrates successful reduction in transcription compared to control across all tissues sampled. Colored dots indicate independent biological replicates. (B) Genome-wide CG methylation level across tissues sampled. Numbers at the top of each bar corresponds to independent biological replicates.

**S3 Fig. Eggs laid in *O. fasciatus* ds-*dnmt1* and control females.** (A) Number of eggs laid by ds-*dnmt1* and control females 8-days post-injection. Dots indicate mean expression level, and error bars indicate standard error of the mean.

**S4 Fig. mCG in *A. mellifera* and *Z. nevadensis* is not associated with transcription.** (A) Gene expression level for deciles of increasing mCG (1–10) and un-methylated genes (UM). Error bars represent 95% confidence interval of the mean. (B) Regression of gene expression against a continuous measure of mCG among all genes with >0 FPKM and weighted mCG. Raw *p* values are provided for each regression, and significance (S) or non-significance (NS) is indicated in brackets following Bonferroni correction.

**S5 Fig. Loss of mCG in *O. fasciatus* ovaries has a limited effect on transcription.** (A) Combinational overlap of genes that are differential CG-methylated and expressed, and similarly CG-methylated and expressed between *O. fasciatus* ds-*dnmt1* and control. Gene groups: Differentially Methylated Gene (DMG), Differentially Expressed Gene (DEG), Similarly Methylated Gene (SMG)/UnMethylated Genes (UMG), and non-Differentially Expressed Gene (non-DEG). More stringent thresholds were used to group genes as CG-methylated or unmethylated (see Materials and Methods).

**S6 Fig. DNA methylation of TEs in *O. fasciatus*.** (A) Levels of mCG across the bodies and 1kb flanking sequence of different annotated TEs in *O. fasciatus*.

**S1 Table. PCR primers used to validate the presence of a single *Dnmt1* ortholog in the *O. fasciatus* genome.**

**S2 Table. Primer sequences for use in producing template DNA for use in the MegaScript transcription kit to generate double stranded RNAs for injection and for quantitative real-time PCR to assess expression levels of *Dnmt1*.**

**S3 Table. Overlap between *O. fasciatus* (control) CG-methylated and unmethylated genes in none and ≥ 1 other insect species used in this study.**

**S4 Table. DNA methylation summary statistics for all species investigated in this study.**

**S5 Table. Output from edgeR v3.20.1 [53] with gene expression (Fragments Per Kilobase of transcript per Million mapped reads [FPKM]) for *O. fasciatus* ovaries ds-*dnmt1* and control biological and technical replicates.**

**S6 Table. Gene expression FPKM for *A. mellifera* queen and drone brains.**

**S7 Table. Gene expression (FPKM) for *Z. nevadensis* female worker at the final instar larva.**

**S8 Table. Output from edgeR v3.20.1 [53] with gene expression (FPKM) for *O. fasciatus* gut, head, and thorax ds-*dnmt1* and control biological replicates.**

**S9 Table. Output General Features File (GFF) from RepeatMasker v4.0.5 (http://www.repeatmasker.org).**

**S10 Table. Significance (*p* value) of TE expression between ds-*dnmt1* and control tissues.**

**S11 Table. Gene Ontology (GO) terms for *O. fasciatus* v1.1 reference genome assembly [26] annotated genes.**

**S12 Table. Significantly enriched GO terms for the intersections between Differentially Methylated Genes (DMG), Similarly Methylated Genes (SMG), Differentially Expressed Genes (DEG), and non-Differentially Expressed Genes (non-DEG).**

**S1 Data. Unaligned and aligned DNA methyltransferase protein sequences in fasta format, and a parenthetical phylogenetic tree in nexus format estimated from the aligned DNA methyltransferase protein sequences using BEAST v2.3.2 [35].**

